# Exercise promotes satellite cell contribution to myofibers in a load-dependent manner

**DOI:** 10.1101/2020.03.19.998211

**Authors:** Evi Masschelein, Gommaar D’Hulst, Joel Zvick, Laura Hinte, Inés Soro-Arnaiz, Tatiane Gorski, Ferdinand von Meyenn, Ori Bar-Nur, Katrien De Bock

**Affiliations:** Department Health Sciences and Technology, Laboratory of Exercise and Health, Swiss Federal Institute of Technology (ETH) Zurich, Zurich, Switzerland; Department Health Sciences and Technology, Laboratory of Regenerative and Movement Biology, Swiss Federal Institute of Technology (ETH) Zurich, Zurich, Switzerland; Department Health Sciences and Technology, Laboratory of Nutrition and Metabolic Epigenetics, Swiss Federal Institute of Technology (ETH) Zurich, Zurich, Switzerland

**Author notes:** contributed equally (Evi Masschelein, Gommaar D’Hulst). Corresponding author: **Katrien De Bock**, Laboratory of Exercise and Health, Department of Health Sciences and Technology (D-HEST), ETH Zürich - Swiss Federal Institute of Technology, ETH Zürich, SLA C7, Schorenstrasse 16, CH-8603 Schwerzenbach, Switzerland.

## Abstract

Satellite cells (SCs) are required for muscle repair following injury and are involved in muscle remodeling upon muscular contractions. Exercise stimulates SC accumulation and myonuclear accretion. To what extent exercise training at different mechanical loads drive SC contribution to myonuclei however is unknown. By performing SC fate tracing experiments, we show that 8-weeks of voluntary wheel running increased SC contribution to myonuclei in mouse plantar flexor muscles in a load-dependent but fiber type-independent manner. Increased SC fusion however was not exclusively linked to muscle hypertrophy as wheel running without external load substantially increased SC fusion in the absence of fiber hypertrophy. Due to nuclear propagation, nuclear fluorescent fate tracing mouse models were inadequate to quantify SC contribution to myonuclei. Ultimately, by performing fate tracing at the DNA level, we show that SC contribution mirrors myonuclear accretion during exercise. Collectively, these findings provide direct evidence that mechanical load during exercise independently promotes SC contribution to existing myofibers.

## Introduction

Muscle tissue has a remarkable ability to adapt its structure and microenvironment in response to diverse conditions such as injury, contractile activity, mechanical overload, and nutrients (Caiozzo, 2002; Chargé and Rudnicki, 2004; Morton et al., 2017). Satellite cells (SCs) are a unique population of stem cells that remain quiescent for the majority of time, but upon activation can proliferate asymmetrically and fuse with myofibers (Tierney and Sacco, 2016). It has been hypothesized that SCs play a supportive role in muscular remodeling by exercise, because SC accumulation as well as myonuclear accretion have been observed during episodes of hypertrophy (Dungan et al., 2019; Fry et al., 2014; Goh et al., 2019; Goh and Millay, 2017). Yet, the quantitative extent of SC contribution to myonuclei was not demonstrated by these studies. Genetic lineage experiments, in which Pax7^+^ cells and their derived fibers are fluorescently labeled, have recently shown that there is a considerable (up to 30%) contribution of SCs to uninjured adult mouse skeletal muscle fibers (Keefe et al., 2015; Pawlikowski et al., 2015). To what extent SCs fuse with existing myofibers during exercise and whether SC fusion mirrors myonuclear accretion under different exercise intensities (with different load) remains to be elucidated.

While there is accumulating data showing SC are required for muscle hypertrophy (Goh et al., 2019; Goh and Millay, 2017) the functional role of SCs in muscle endurance adaptations is much less understood (Abreu et al., 2017). Running exercise increases the number of myofibers containing an elevated number of SCs (Cisterna et al., 2016; Shefer et al., 2010) and SC number is directly proportional to running performance (Kurosaka et al., 2009). Nevertheless, SC number but not myonuclear number increased in several rat hindlimb muscles after a period of free wheel running (Smith and Merry, 2012), indicating that SC accumulation and myonuclear accretion are not necessarily linked. Moreover, no defects in intrinsic adaptations to endurance exercise (eg. muscle vascularization, fiber type shift or succinate dehydrogenase activity) were reported when SCs were ablated using the *Pax7^CreERT2-DTA^* model or X-ray irridation, arguing against the necessity of SCs for adaptations to endurance exercise (Jackson et al., 2015; Li et al., 2006). Consequently, it is currently unknown whether exercise training aimed to evoke endurance adaptions, but not muscle hypertrophy, can stimulate SC contribution to myonuclei.

An excellent model to study the involvement of SCs in muscle adaptations to exercise is voluntary wheel running. Mice have a strong intrinsic drive to run large distances in their active phase (De Bono et al., 2005), which keeps stress and mouse handling at a minimum. Furthermore, by applying resistance to the running wheel, muscle hypertrophy can be induced without substantial muscle damage (Cornelison, 2018; Legerlotz et al., 2008; Mccarthy et al., 2017; Murach et al., 2018; O’Connor et al., 2007; Soffe et al., 2016; White et al., 2016) and reducing wheel resistance (free running) leads to favorable endurance adaptations without pronounced effects on muscle hypertrophy (Jackson et al., 2015; Manzanares et al., 2019). Thus, using this exercise model in combination with genetic lineage tracing experiments of SCs, we aimed to evaluate whether SC contribution to myonuclei is mediated by increasing load during exercise training and whether this manifests in SC-mediated myonuclear accretion.

## Methods

### Animals

All mice were housed in individually ventilated cages (3-4 littermates per cage) at standard housing conditions (22°C, 12 h light/dark cycle), with ad libitum access to chow (KlibaNafag, diet #3436 and diet #3437) and water. Health status of all mouse lines was regularly monitored according to FELASA guidelines. All mouse lines were maintained on a C57BL/6J background: *Pax7^CreERT2^* (Jackson labs stock no: 017763), *Rosa^mTmG^* (Jackson labs stock no: 007576) and *Rosa^CAG-LSL-ntdTomato^* (Ai75D, Jackson labs stock no: 025106). *Rosa^mTmG^* and *Rosa^CAG-LSL-ntdTomato^* mice were intercrossed with *Pax7^CreERT2^* mice to generate *Pax7^CreERT2/+^*; *Rosa^mTmG/+^* (*Pax7^mTmG/+^*) and *Pax7^CreERT2/+^*; *Rosa^CAG-LSL-ntdTomato/+^* (*Pax7^nTom/+^*). All mice used in this study were female. At the age of 15 weeks, mice received tamoxifen via intraperitoneal injection at a dose of 1.0 mg/day for 5 days to induce Cre-mediated recombination and a washout period of 7 days was allowed before experiments were initiated.

### Experimental procedures

All animal procedures were approved by the Veterinary office of the Canton of Zürich (license nr. ZH254-16). Mice were individually housed in open cages equipped with a running wheel device (TSE Systems, Bad Homburg vor der Höhe, Germany) for the duration of the intervention. Non-running control mice were also singly housed in cages of equal dimensions, but without running wheels. The running wheel device continuously records wheel movements out of which total distance (km), speed (m.s^−1^), number of running bouts and resistance on the wheel (N) were extracted. Additionally, to increase the force needed to rotate the wheel, resistance (0-100%) can be added. To calculate total external work we used the equation W = Pt where W is work, P is power, and t time. To calculate the Power of the wheel at each braking resistance we used the equation P = 2π × f × M where f is the angular frequency of the wheel, and M is the torque at a given braking resistance. TSE Systems provided a torque braking resistance curve. Mice were randomized in three groups: sedentary controls (no running), voluntary low load endurance wheel running where there is no extra resistance on the wheel (VRun) and voluntary high load resistance wheel running where extra resistance is placed on the wheel (VResRun). A representation of the study design is shown in Figure 1A. Prior to the 8 weeks of voluntary wheel running, both groups VRun and VResRun were familiarized for 5 days to the running wheel without extra resistance. After familiarization, the VResRun group was subjected to a progressive weekly increase in resistance on the wheel: The load on the wheel was 50% in week 1, 60% in week 2, 70% in week 3, 72% in week 4, and 74% from week 5 to 8. The minimal distance which mice needed to run was set at 2km per night. In case a mouse did not reach 2km for 2 consecutive nights, resistance was slightly reduced for 3 nights after which it was raised again. All mice ran at 74% resistance during the last 2 weeks.

**Figure 1.**
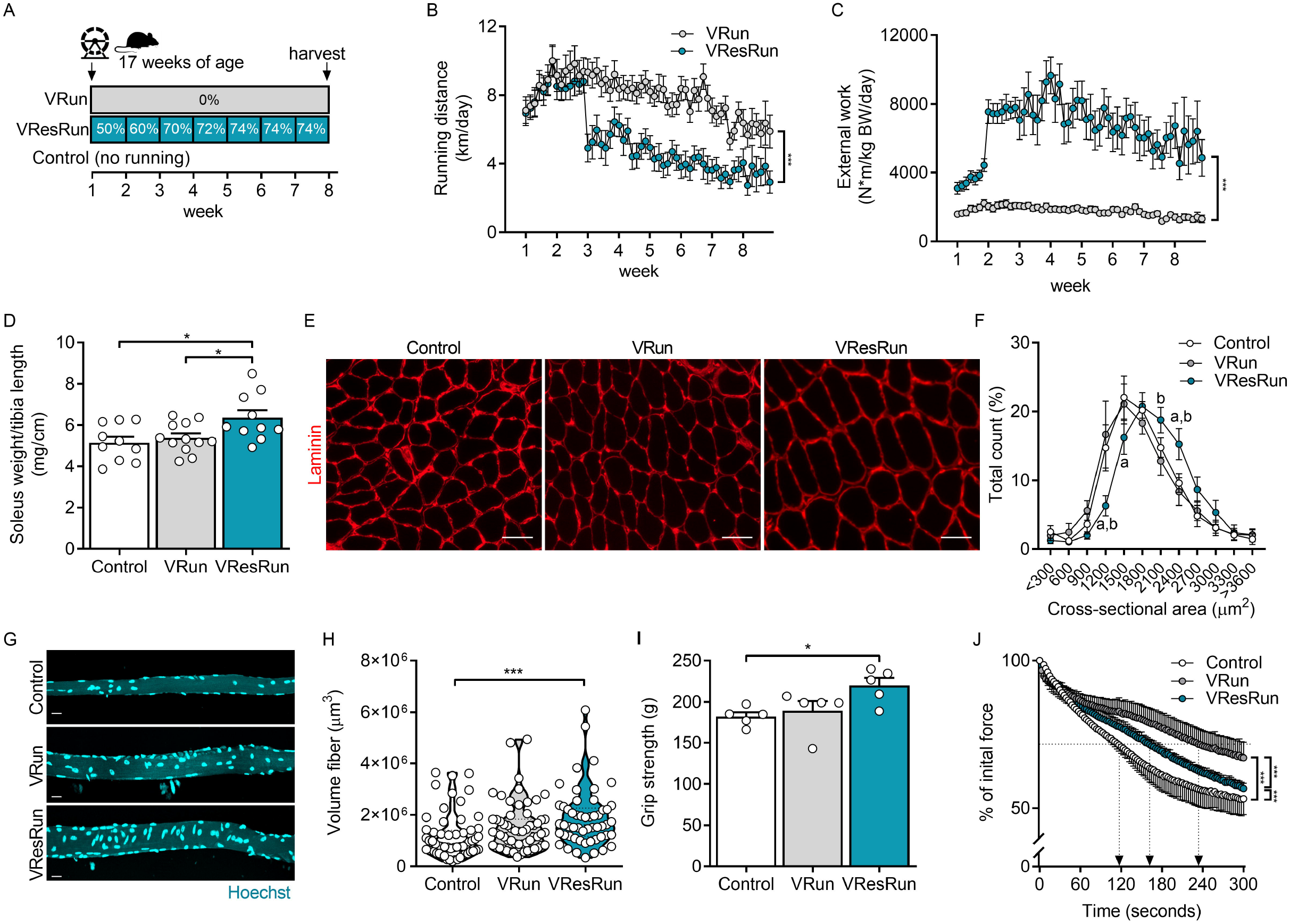
Running pattern and muscle hypertrophy is load-dependent. (A) Experimental set-up. Average running distances (B) and average external work (C) per night throughout the exercise protocol (n = 16 mice per group). (D) Soleus weight normalized to tibia length. Representative cross-sections stained for Laminin (red) (E) and quantification (n = 7-8 mice per group) of fiber cross-sectional area distribution in soleus muscle (F). Representative pictures of single myofibers isolated from soleus (G) and quantification of fiber volume of 11-16 single myofibers per mouse (n = 4 mice per group). (H) Muscle force production by grip strength (I) and time-to-fatigue in *ex vivo* stimulated muscle (J). Statistics: one-way ANOVA test with Tukey correction for multiple comparisons (F,H,I) or two-way ANOVA test with a Bonferroni post hoc test (B,C,D,J). (*p < 0.05; **p < 0.01; ***p < 0.001) (a, p < 0.05 VResRun compared to Control; b, p < 0.05 VResRun compared to VRun). Each dot represents a single mouse (D,H,I) or muscle fiber (H). Bar graphs (D,I) and line graphs (B,C,F,J) represent mean ± SEM (error bars). Violin plot (H). Scale bars, 50 μm (E) and 25 μm (G).

### Sample Collection

Tissue collection was performed 24 h fter the last exercise session after a 4 h fast. Mice were anaesthetized using Ketamine (80-100 mg/kg), Xylazine (10-15 mg/kg) and Acepromazine (2-5 mg/kg) via intraperitoneal injection 5 min before sacrifice. The depth of anesthesia was confirmed by testing pedal withdrawal reflex. Subsequently, the m. soleus (SOL), m. plantaris (PLT), m. gastrocnemius (GAS), m. tibialis anterior (TA), and m.extensor digitorum longus (EDL) were harvested, weighed and frozen in OCT embedding matrix (CellPath) in liquid nitrogen-cooled isopentane for histochemical analysis or fixed in 4% PFA for single myofiber isolation. Tibia length was assessed by a digital caliper. After sample collection, animals were euthanized and major bleeding was induced to confirm death.

### Immunofluorescence

Frozen sections (10μm) of muscle embedded in OCT of mid-belly level were made using a cryostat (Leica CM 1950) and collected on Superfrost Ultra Plus slides (Thermo Scientific, Zug, Switzerland). The following antibodies were used: anti-MyHC-I (1:50 dilution, BA-F8 from hybridoma, Iowa City, IA, USA), anti-MyHC-IIa (1:200 dilution, SC-71 from hybridoma), anti-MyHC-IIb (1:100 dilution, BF-F3 from hybridoma), anti-laminin (1:200 dilution, PA1-16730, ThermoFisher), anti-PCM1 (1:1000 dilution, HPA023370, Sigma, Buchs, Switzerland), Alexa Fluor 488 goat antimouse IgG2B (1:250 dilution, ThermoFisher), Alexa Fluor 350 goat anti-mouse IgG1 (1:250 dilution, A21120, ThermoFisher), Alexa Fluor 568 goat anti-mouse IgM (1:250 dilution, A-21043, ThermoFisher), Alexa Fluor 647 goat anti-rabbit IgG (1:250 dilution, A-21244, ThermoFisher). Alexa Fluor 568 goat anti-mouse IgG1 (1:400 dilution, A-21124, ThermoFisher), Alexa Fluor 647 conjugated wheat germ agglutinin (WGA, 1:50 dilution, W32466, ThermoFisher).

Immunohistochemical muscle fiber staining was conducted as previously described (Bergmeister et al., 2016). In short, sections were dried and washed for 5 min in PBS supplemented with 0.05% Triton-X-100 (PBST) and subsequently blocked for 60 min in PBST + 10% goat serum (16210064, ThermoFisher). Afterwards a primary antibody cocktail diluted in PBST + 10% goat serum was applied for 120 min against MyHC-I, MyHC-IIa, MyHC-IIb and laminin. After washing 3 times for 5 min, a secondary antibody cocktail, diluted in PBST + 10% goat serum, was applied for 60 min. Slides were mounted after a 3 x 5 min wash and sealed with glass cover slips.

For myonuclear staining, samples were stained with anti-PCM1 as previously described (Winje et al., 2018). Slides were pre-incubated in 2% BSA in PBS. Sections were stained with a rabbit polyclonal antibody against PCM1 in staining solution (5% BSA in PBS, 0.2% Triton-X-100) overnight at 4°C. The next day, the sections were washed three times for 5 minutes in PBS and stained with an antirabbit secondary antibody in 2% BSA in PBS for 1 h. Sections were washed three times for 10 minutes with PBS. For detection of the fiber boundaries, the sections were stained with WGA in PBS for 30min. Nuclei were co-stained using Hoechst dye 33342 (dilution 1:5000, H3570, ThermoFisher).

Pax7 staining was performed as previously described (Bar-Nur et al., 2018). Slides were cross-linked with 4% PFA, washed with 1xPBS, incubated for 30 minutes with 2% BSA, dissolved in PBS and 0.1% Triton-X-100, followed by 15 minutes incubation with 10% donkey serum (017-000-121, Jackson ImmunoResearch, Cambridgeshire, UK) diluted in PBS. Sections were then incubated for 1h in primary antibodies against Pax7 and laminin at room temperature followed by PBS rinse (twice) and incubation in secondary antibodies (1h at room temperature). Nuclei were co-stained using Hoechst dye 33342 (dilution 1:5000, H3570, ThermoFisher).

Images from sections were captured at 10x using an epifluorescent microscope (Zeiss Axio observer Z.1, Zeiss, Oberkochen, Germany). Composite images were stitched together using the Tiles module in the ZEN 2011 imaging software (Zeiss). All images were captured at the same exposure time.

Satellite cell contribution to myofibers in *Pax7^mTmG/+^* mice was quantified based on a previously described method of myofiber masking (Tierney et al., 2018). Images were processed using an Image-J plug-in to define myofiber cross-sectional area (CSA) (Noirez et al., 2006), followed by a manual refinement of the CSA outlines. An overlay of those outlines on the corresponding mGFP image was performed, after which the data was compiled in Excel and a threshold was set for the mean gray value of mGFP. A frozen section isolated from a vehicle treated mouse (negative control sample) was used to determine the threshold and to ensure the elimination of autofluorescence. mGFP images were overlaid with the muscle fiber type staining to identify mGFP^+^ oxidative (MyHCI) and glycolytic (MyHCIIa, MyHCIIb and MyHCIIx) fibers.

### Single myofiber isolation

For single myofiber analysis, the SOL from the *Pax7^nTom/+^* mice was harvested, fixed in 4% paraformaldehyde/PBS for 1h at 4°C, and transferred to PBS containing 2% horse serum. To release individual myofibers, single fibers were mechanically teased apart, strained, and washed with PBS. Isolated single myofibers were stained with Hoechst for nuclear visualization and gently mounted on a glass coverslip. Images were captured at 10x using a confocal microscope (Olympus olympus fluoview FV 3000, Olympus, Hamburg, Germany). Eleven to sixteen fibers per mouse from random areas from the muscle were analyzed. Total number of nTom^+^ nuclei and Hoechst^+^ nuclei along 500 μm fiber length were quantified using a spot detection algorithm developed in Imaris (Bitplane). Myofiber volume was approximated as the volume of a cylinder using the average radius measured along the fiber length in Imaris. Fibers isolated from vehicle-treated mice were used to set a threshold to ensure the elimination of autofluorescence.

### Cell culture

#### Reagents

Freshly isolated primary myoblasts were cultured in growth medium, which contained a 1:1 ratio DMEM (ThermoFisher Scientific, 12320032) and Ham’s F-10 nutrient mix (ThermoFisher Scientific, 22390058) supplemented 20% FBS (Thermo Scientific, 10270106) and 10 ng/ml basic-FGF (ThermoFisher Scientific, PHG0266). Cells were differentiated in low-serum differentiation medium (Thermo Scientific, 41965039), supplemented with 2% horse serum (Thermo Scientific, 16050-122). Cells were routinely cultured in 21% O_2_ and 5% CO_2_. Every other day, differentiation medium was changed until myotubes reached full differentiation. All media were supplemented with 100 units/ml penicillin and 100 μg/ml streptomycin.

#### Isolation of SCs/myoblasts

Isolation of SCs/myoblasts was done as previously described (D’Hulst et al., 2020). Briefly, muscle tissue was digested in Hank’s Balanced Salt Solution (HBSS, Thermo Fisher, 24020117) supplemented with 1.5% bovine serum albumin (BSA) and 2 mg/mL collagenase type II (ThermoFisher Scientific, 17101015) for 1 h at 37 °C. After centrifugation, the cell pellet was then filtered using 100 and 40 μm cell strainers and a heterogeneous cell population was purified by FACS sorting or by serial preplating. For FACS, SCs were sorted based on positive alpha 7-integrin (1:100, R&D Systems FAB3518P) and absence of Sca1 (1:1000, Bio-Legend 122511), CD31 (1:1000, Bio-Legend, 102413), and CD45 (1:1000, Bio-Legend 103121).

#### Generation of nuclear H2B-GFP (H2B-nGFP) myoblasts

Lentivirus was generated with PEG-it Virus Precipitation Solution (System Biosciences, Palo Alto, CA, USA no LV810A-1) using a LV H2B-GFP plasmid (Addgene no 25999) in HEK 293 cells. Myoblasts were transfected with 20μl of concentrated lentivirus for 24h using growth medium supplemented with 4μg/ml polybrene transfection reagent (TR1003-G, Sigma-Aldrich).

#### Co-culturing experiments

nTom^+^ myoblasts, together with committed WT myoblasts were co-cultured at 1:4 ratio (nTom^+^/WT). nTom^+^ myoblasts were co-cultured with myoblasts stably expressing H2B-nGFP at a ratio of 1:1 ratio (nTom^+^/H2B-nGFP^+^). Two hours after seeding, growth medium was changed to differentiation medium. Every other day, differentiation medium was changed until myotubes reached full differentiation and immunofluorescent images were taken every 24h via an epifluorescent microscope, equiped with a cell incubator (Nikon eclipse T*i*2, Nikon, Amstelveel, Netherlands). Fluorescent pictures were captured every 15 min to generate a time-lapse video (Windows movie maker, Windows, Redmond, WA, US).

### SC fate tracing at the DNA level using RT-PCR for Cre-mediated recombination in bulk muscle and sorted myonuclei

To perform SC fate tracing at the DNA level, genomic DNA was isolated from muscle or myonuclei using QIAamp DNA Micro and Mini Kit (Qiagen, Hilden, Germany). Myonuclei were isolated as previously described (Habib et al., 2017). Nuclei were isolated with EZ PREP buffer (Sigma, NUC-101). Frozen soleus muscles (10 mg) were homogenized using a glass dounce tissue grinder in 2 ml of ice-cold EZ PREP and incubated on ice for 5 min, with an additional 2 ml of ice-cold EZ PREP and filtered through a 100 μm cell strainer. Nuclei were centrifuged at 500 × g for 5 min at 4 °C, washed with 4 ml ice-cold EZ PREP and incubated on ice for 5 min. After centrifugation, the nuclei were stained for 45 min with an antibody against PCM-1 (1:1500, HPA023370, Sigma), followed by an Alexa647-anti-rabbit secondary antibody for 30min (1:1500) and Hoechst 33342 (1:5000), and filtered through a 35 μm cell strainer. Hoechst^+^ and Alexa647^+^ myonuclei were directly sorted into RLTplus buffer (Qiagen). A final amount of 10,000-20,000 nuclei was used for genomic DNA isolation. A SYBR Green-based master mix (ThermoFisher Scientific, A25778) was applied for RT-PCR analysis. Recombination rates were calculated from the relative expression of recombined levels normalized to an internal control. The delta–delta CT method was used to normalize the data. The percentage recombination was calculated relative to positive controls. To generate a positive control, we crossed *HSA^CreERT2^* mice (McCarthy et al., 2012) with *Rosa^mTmG^* mice to generate *HSA^CreERT2/+^;Rosa^mTmG/+^* mice. In these mice, tamoxifen injection leads to the excision of *mTomato* from the DNA in myonuclei, and the subsequent expression of *mGFP*. Additionally, we sorted SCs from *Pax7^nTom/+^* mice. We assumed that recombination was 100% in myonuclei from *HSA^CreERT2/+^;Rosa^mTmG/+^* mice as well as in SCs from *Pax7^nTom/+^* mice. The following primers were used to detect recombination (see also Figure 5A and 5D): *Rosa^mTmG^*, recombined amplicon: Fw GGGCTCGACACTAGTGAACC Rv GGTGATGATCCGGAACCCTT, Internal control: Fw AGCGAACGGACAGGAGAATG, Rv ACTTGTGGCCGTTTACGTCG, *Rosa^CAG-LsL-ntdTomato^*, recombined amplicon: Fw TTATTGTGCTGTCTCATCATTTTGG Rv CTTCGCGAGACCGGTAATCT, Internal control: Fw CTGTTCCTGTACGGCATGG Rv GGCATTAAAGCAGCGTATCC.

**Figure 5.**
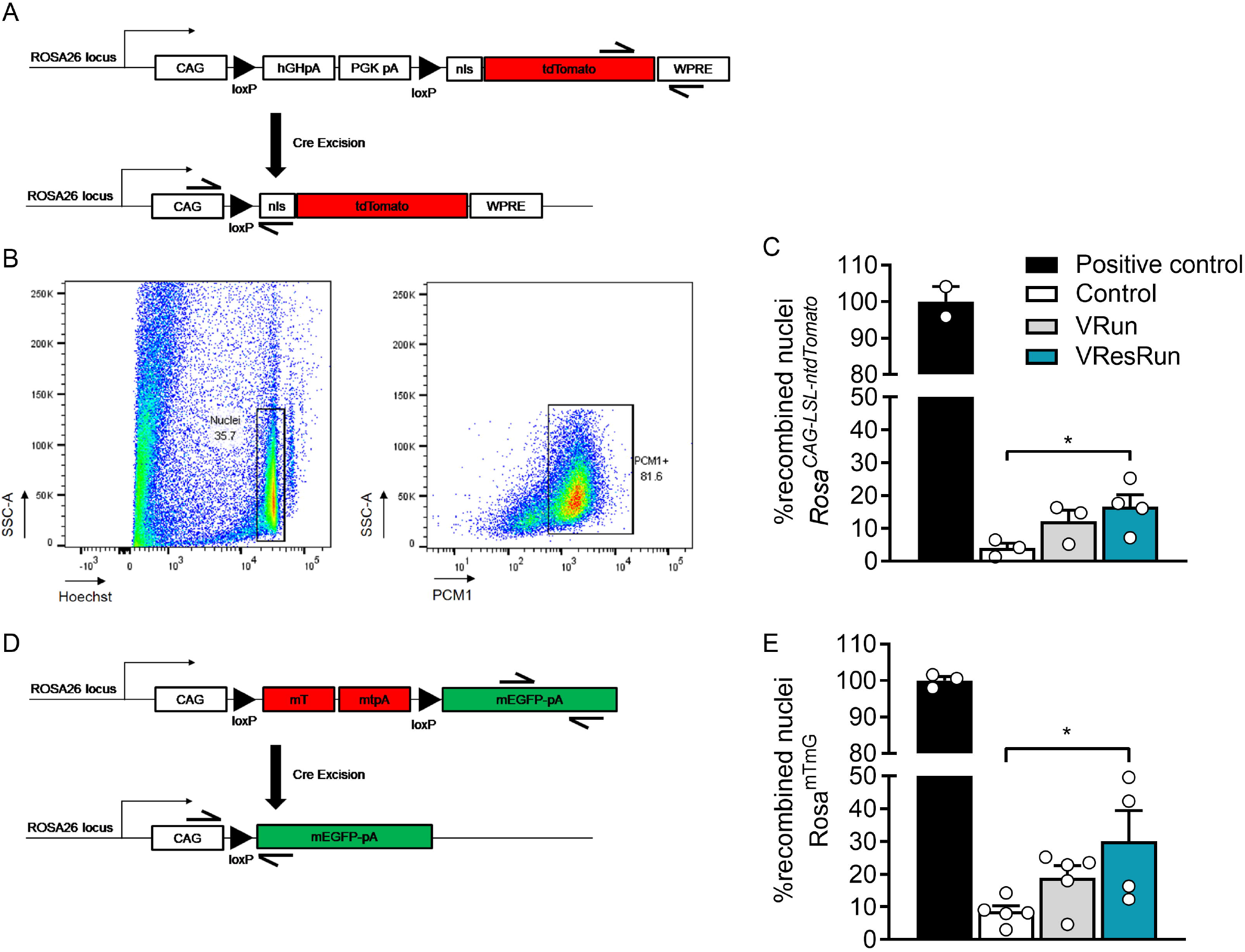
Myonuclear-specific Cre-mediated recombination. (A) Schematic diagram of the *Rosa^CAG-lsl-ntdTomato^* construct before and after Cre-mediated recombination. Representation of primers used for quantitative RT-PCR analysis showing amplicon of recombinant DNA and amplicon of internal control for normalization. (B) Representative flow cytometry plots depicting the gating strategy for sorting of myonuclei. Isolated nuclei were analyzed by side scatter (SSC) and Hoechst. Hoechst^+^PCM1^+^ single nuclei were sorted for further analysis. (C) RT-PCR performed on sorted Hoechst^+^PCM1^+^ nuclei. DNA isolated from satellite cells from *Pax7^CreERT2/+^; Rosa^CAG-lsl-ntdTomato/+^* mice with the presence of one recombined allele in each satellite cell served as a 100% reference (positive control). (D) Schematic diagram of the *Rosa^mTmG^* construct before and after Cre-mediated recombination. Representation of primers used for quantitative RT-PCR analysis showing amplicon of recombinant DNA and amplicon of internal control for normalization. (E) RT-PCR performed on genomic DNA isolated from bulk muscle. DNA isolated from tamoxifen injected *HSA^iCre/+^;Rosa^mTmG/+^* mice with the presence of one recombined allele in each myonucleus, served as a 100% reference (positive control). Recombination rates were calculated from the relative expression of recombined levels normalized for internal control. Statistics: one-way ANOVA test with Tukey correction for multiple comparisons (*p < 0.05; **p < 0.01; ***p < 0.001). Each dot represents a single mouse. Bar graphs represent mean ± SEM (error bars).

### Grip strength

Grip strength of the limbs was measured using a force tension apparatus (grip strength meter, Bioseb, Vitrolles, France) after 8 weeks of voluntary wheel running or no running. Mice were lifted by the tail and were made to hold a metal grid with all limbs. Total-limb maximal grip strength was registered in gram during three consecutive attempts, and the result was set as the average of the attempts.

### Ex vivo muscle force assay

The SOL was carefully dissected and placed in Krebs–Henseleit buffer (120 mM NaCl, 4.8 mM KCl, 25 mM NaHCO3, 2.5 mM CaCl2, 1.2 mM KH2PO4, 2 mM MgSO4) supplemented with 25 mM glucose at 37°C and bubbled with 95% O_2_–5% CO_2_ to stabilize the pH at 7.4. The distal tendon was securely connected to a fixed bottom plate, and the proximal tendon to the arm of a servomotor (800A in vitro muscle apparatus, Aurora Scientific). Muscle length was adjusted until a single stimulus pulse elicited maximum force during a twitch (optimal length, Lo) under isometric conditions. After 15min of equilibration, a fatigue protocol was started. To fatigue the muscle, tetanic contractions of 60 Hz were produced, once every 2 s, for a total of 300 s. Time-to-fatigue at 75%, an assessment for the time it takes for a muscle to fall to 75% of its initial force, was calculated for every mouse.

### Statistical analysis

Statistical significance was determined by a one-way ANOVA with Tukey correction for multiple comparisons or two-way ANOVA with Bonferroni correction with Graphpad Prism 8.2. Data are presented as means ± SEM, and values of p < 0.05 were considered statistically significant. The degree of significance is depicted as follows: *p < 0.05, **p < 0.01, ***p < 0.001.

## Results

### Running pattern, hypertrophy and muscle performance in VRun vs. VResRun

To study how increased load during exercise affects contribution of SCs to myofibers, we used 8 weeks of voluntary low load - endurance type - running (VRun) and voluntary high load - resistance type - running (VResRun) as exercise models (Figure 1A). A progressive increase in load reduced running distance (Figure 1B), running speed and total daily running time without affecting the number of running bouts from week 3 onwards (Figure 1-figure supplement 1A-C). In contrast, external work was higher over the 8 week time period (Figure 1C). In agreement with previous observations (Konhilas et al., 2005; Legerlotz et al., 2008; White et al., 2016), extra resistance on the wheel caused an increase in the weight of m. soleus (SOL), while unloaded running did not change SOL weight (Figure 1D). Body weight, lean mass and fat mass were not affected by exercise (Figure 1-table supplement 1). Further analysis on cross-sectional area (CSA) confirmed that VResRun, but not VRun, induced fiber hypertrophy in SOL (Figure 1E,F). This was exclusively caused by an increase in the CSA of MyHCIIa muscle fibers, while MyHCI fiber CSA remained unaffected (Figure 1-figure supplement 2A-E). Finally, in accordance with the data on whole muscle, VResRun increased the volume of isolated single myofibers from SOL, while VRun did not affect myofiber volume (Figure 1G,H).

Despite unchanged muscle weight in PLT (Figure 1-table supplement 1), we measured increased fiber CSA in PLT upon VResRun as well as VRun. CSA of MyHCIIa fibers increased both after VRun and VResRun compared to control, whereas MyHCIIb-IIx fiber CSA only increased after VResRun, (Figure 1-figure supplement 2F-J). The discrepancy between muscle mass and CSA in the PLT could be due to difficulties in anatomically dissecting the muscle (Egner et al., 2016). Muscle mass of m. gastrocnemius (GAS), m. tibialis anterior (TA) and m. extensor digitorum longus (EDL) remained unaffected by VRun and VResRun (Figure 1-table supplement 1). In addition, no change in fiber CSA was found for GAS, TA and EDL (Figure1-figure supplement 2K,L,M).

Finally, to analyze whether the running-induced increases in CSA also resulted in increased muscle force production, we tested muscle strength by grip strength experiments. After 8 weeks, grip strength was increased in VResRun (Figure 1I). Time-to-fatigue at 75%, an assessment for the time it takes for a muscle to fall to 75% of its initial force, was higher after VRun and VResRun compared to control (Figure 1J). Collectively, these data confirm that VRe

sRun, but not VRun, induces an increase in muscle mass in both SOL and PLT, which was induced by hypertrophy of the glycolytic muscle fibers.

### Exercise promotes SC contribution to myofibers in a load-dependent manner

To investigate whether voluntary wheel running affects SC contribution to various hind limb muscles, and whether this is load-dependent, we genetically labeled SCs and performed fate-tracing experiments into myonuclei. To do so, we used a mouse model which was previously used to study SC involvement in muscle regeneration (Murphy et al., 2014) and muscle homeostasis (Keefe et al, 2015). Briefly, we intercrossed *Pax7^CreERT2/CreERT2^* mice, which efficiently induces recombination in SCs, with mice carrying a *Rosa^mTmG/mTmG^* reporter (Muzumdar et al., 2007). The resulting *Pax7^CreERT2/+;^ Rosa^mTmG/+^* mice (*Pax7^mTmG/+^*) ubiquitously express membrane-bound Tomato (*mTomato*), but following tamoxifen injection, Cre-mediated recombination results in the excision of *mTomato* and expression of membrane-bound GFP (*mGFP*) in SCs. In these mice, the presence of GFP^+^ myofibers indicates that at least one SC fused with those myofibers. In our experiments, we labeled SCs by 5 consecutive daily IP tamoxifen injections and initiated the exercise protocol 1 week later. Subsequently, hindlimb muscles were harvested after 8 weeks of VRun or VResRun or no running (control) (Figure 2A) and the contribution of SCs to myofibers was evaluated. Importantly, we opted to use 15 week old mice for all our experiments since this is an age where postnatal muscle growth has ceased and SC contribution to myofibers is restricted to maintenance of muscle homeostasis (Bruusgaard et al., 2006; Murach et al., 2017; Pallafacchina et al., 2013; Pawlikowski et al., 2015; White et al., 2010). Indeed, in agreement with previous work (Keefe et al., 2015; Pawlikowski et al., 2015), we found that in the non-exercising control mice, 5-20% of all myofibers were GFP^+^, with the highest contribution of SCs in the SOL, confirming that SC fusion occurs in hindlimb muscles of sedentary mice (Figure 2B,C). Interestingly, running increased the SC contribution to SOL, PLT and GAS, whereas we did not observe a clear increase in SC contribution to fibers upon exercise in TA, nor EDL (Figure 2B,C), confirming previous observations from our group that wheel running specifically recruits the hindlimb plantar flexors (D’Hulst et al., 2019). Moreover, SC contribution to the SOL was load-dependent, as VResRun further increased the number of GFP^+^ myofibers compared to VRun alone, resulting in 80% GFP^+^ myofibers vs. 60% in VRun condition. In PLT and GAS, we also observed a 38% increase in GFP^+^ myofibers upon VResRun, while the increase by VRun did not reach statistical significance (Figure 2B,C)

**Figure 2.**
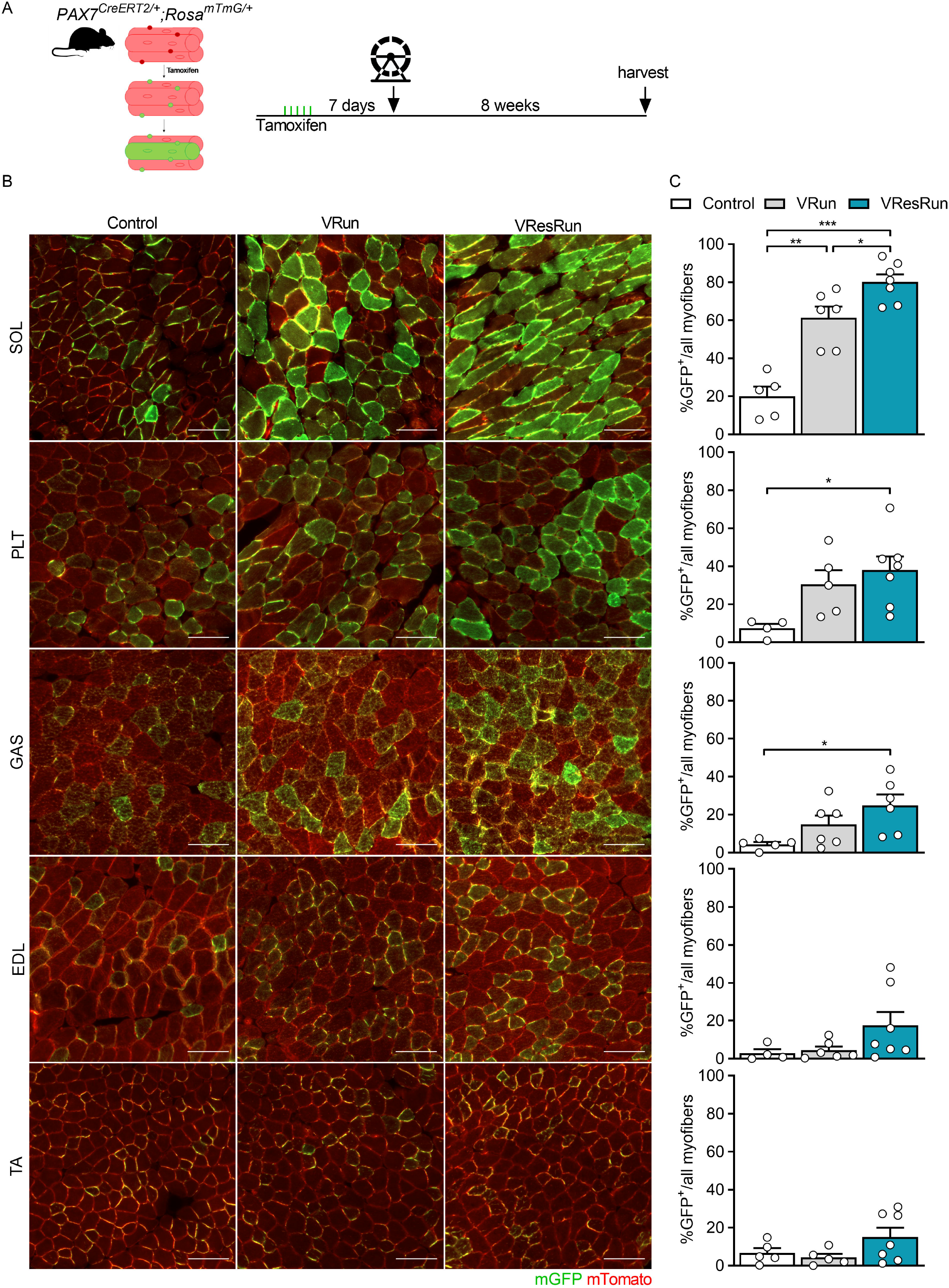
Exercise promotes SC contribution to myofibers in a load-dependent manner. (A) *Pax7^CreERT2^* mice were crossed with *Rosa^mTmG^* mice to genetically trace SC fusion to myofibers. Upon tamoxifen treatment SCs will exoress mGFP. mGFP^+^ myfibers indicate fusion of SCs. (B) Representative cross-sections of soleus (SOL), plantaris (PLT), gastrocnemius (GAS), tibialis anterior (TA) and extensor digitorum longus (EDL) muscle showing GFP^+^ myofibres to which SCs contributed. (C) Quantification of GFP^+^ myofibers. Statistics: one-way ANOVA test with Tukey correction for multiple comparisons (*p < 0.05; **p < 0.01; ***p < 0.001). Each dot represents a single mouse. Bar graphs represent mean ± SEM (error bars). Scale bar, 100 μm.

To investigate whether the increase in SC fusion by exercise was fiber type dependent, we made sequential cryosections of SOL, overlaid the GFP signal with fiber type immunostainings and assessed the relative amount of GFP^+^ fibers to MyHC type (Figure 2-figure supplement 3A). VResRun induced the largest increase in GFP^+^ fibers resulting in ~80% of all MyHCI and MyHCIIa fibers becoming GFP positive, while this was only ~60% in VRun, indicating that increasing load promoted SC contribution to myofibers but that there was no fiber type preference (Figure 2-figure supplement 3B,C). Moreover, in agreement with previous data (Yan et al., 2011), both running modalities led to a shift towards a more oxidative fiber type (MyHCI) with the largest shift in the VRun group (Figure 2-figure supplement 3D). Thus, because SC fusion occurred to the same extent in MyHCI and MyHCIIa, relative GFP contribution after exercise mirrored the shift in fiber type (Figure 2-figure supplement 3E). Alltogether, we show that exercise drives SC conribution to muscle fibers in a load-dependent manner, but independent of muscle fiber type.

### Nuclear propagation renders nuclear reporter mice inadequate to assess SC contribution to myonuclei

Our data from the *Pax7^mTmG/+^* mice indicated that exercise strongly increases SC contribution to myofibers. Nevertheless, this model does not allow us to assess the exact number of SCs that fuse with the myofiber since the GFP signal diffuses throughout the sarcolemma, nor does it provide information on myonuclear accretion upon exercise training. Therefore, we generated a nuclear reporter mouse by crossing the aforementioned *Pax7^CreERT2^* with a *Rosa^CAG-LSL-ntdTomato^* mouse. The *Rosa^CAG-LSL-ntdTomato^* mouse expresses a loxP-flanked stop codon before the nuclear tdTomato fluorescent red (nTom) reporter. Upon tamoxifen injection, the nuclear stop codon is excised to indelibly label Pax7^+^ SCs and their derivative nuclei with nTom (Figure 3A). We used the *Pax7^CreERT2/+^;Rosa^CAG-LSL-ntdTomato/+^* (*Pax7^nTom/+^*) mice to assess myonuclear accretion and SC contribution to myonuclei by isolating single muscle fibers from SOL (Figure 3A) after 8 weeks of exercise training.

**Figure 3.**
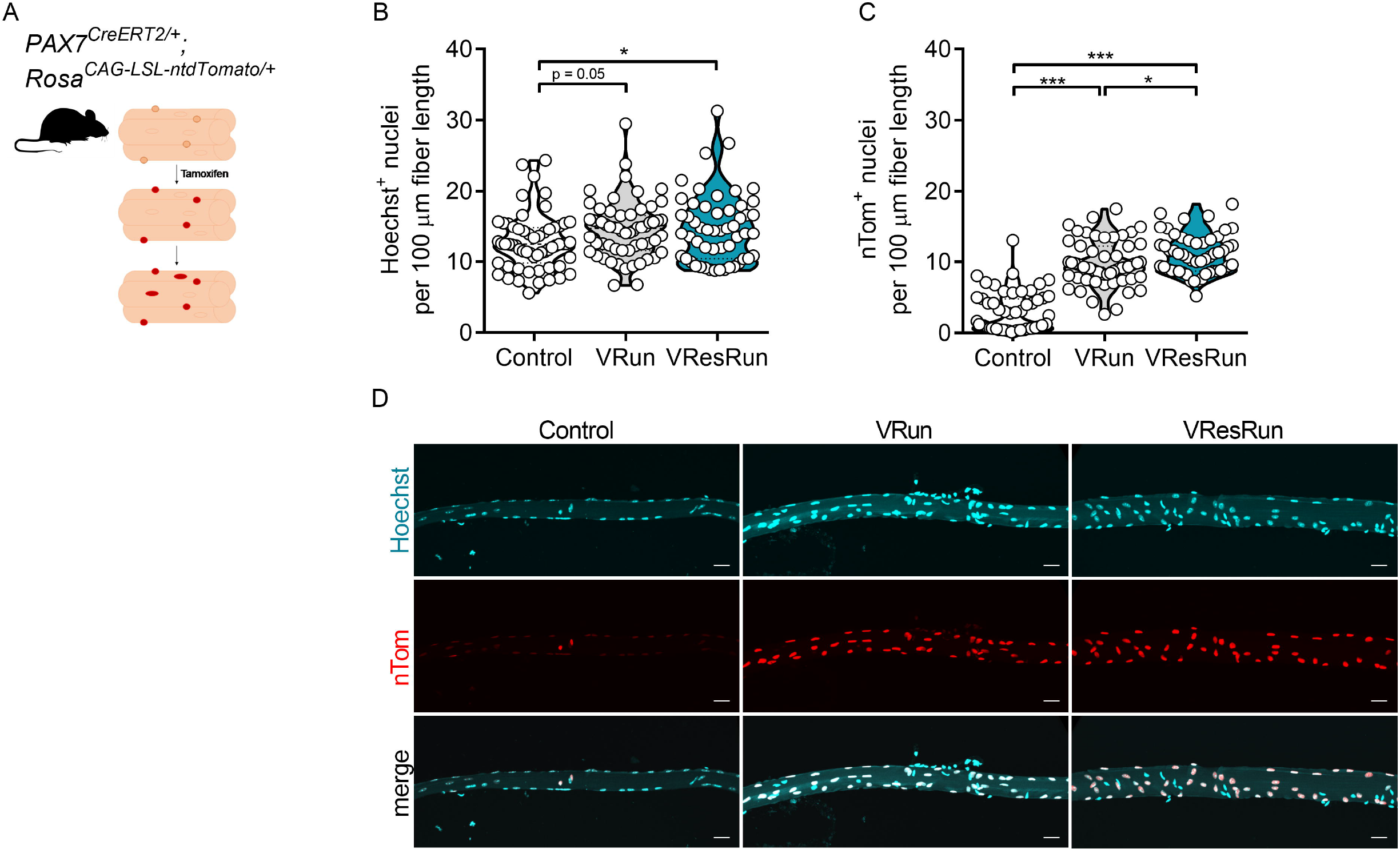
Nuclear reporter mice show high SC contribution during exercise. (A) *Pax7^CreERT2^* mice were crossed with *Rosa^CAG-LSL-ntdTomato^* mice to genetically trace SC-derived nuclei. Upon tamoxifen treatment, SCs and SC-derived myonuclei will express nTomato (nTom). Quantification of Hoechst^+^ (B) and nTom^+^ nuclei (C) per 100μm fiber length (11-16 single myofibers per mouse, n = 4 mice per group). (D) Represented pictures of single myofibers isolated from soleus. Statistics: one-way ANOVA test with Tukey correction for multiple comparisons (*p < 0.05; **p < 0.01; ***p < 0.001). Violin plot is shown (B,C). Each dot represents a single mouse. Scale bar, 25 μm.

VResRun, and to a lesser extent VRun, increased the number of Hoechst^+^ nuclei on single fibers compared to control by respectively 16 and 14% (p<0.05 and p=0.05, Figure 3BC). Myonuclear specific PCM1 staining on cryosections corroborated these findings (Figure 3-figure supplement 4A), indicating that myonuclear accretion after exercise is load-dependent. Intriguingly, compared to myonuclear (Hoechst^+^) accretion, we observed much larger increase in nTom^+^ nuclei after both VRun (+1.8 Hoechst^+^ nuclei vs. +7.0 nTom^+^ nuclei per 100 μm fiber length) and VResRun (+2.0 Hoechst^+^ nuclei vs. +8.3 nTom^+^ nuclei per 100 μm fiber length) (Figure 3B,C,D), suggesting 1) high turnover of myonuclei during exercise or 2) nuclear reporter mice are inadequate to measure SC fusion in myocytes due to nuclear travelling of nTom, a process recently termed ‘propagation’ by Taylor-Weiner et al. (Taylor-weiner et al., 2020). To investigate the latter hypothesis, we performed a series of in vitro coculturing experiments. In a first experiment, we cultured nTom^+^ myoblasts together with myoblasts which stably expressed GFP in the nucleus (H2B-nGFP) at a 1:1 ratio and differentiated them to myotubes for 144h (Figure 4A). In this experiment, protein propagation between nuclei in multinucleated myotubes would lead to nTom+/H2B-nGFP^+^ myonuclei. Time-course imaging revealed that once H2B-nGFP^+^ and nTom^+^ myoblast started to fuse, double positive nuclei (yellow) became apparent, resulting in 45% of all nuclei that turned double positive for H2B-nGFP and nTom at full differentiation (Figure 4B,C), showing that there is reporter protein propagation. Since only limited number of SCs contribute to myonuclei in myofibers *in vivo*, we performed a second experiment, where we cultured nTom^+^ myoblasts together with committed myocytes which do not express a reporter protein (WT) at a ratio of 1:4 (nTom^+^/WT) and further differentiated them to myotubes for 144h (Figure 4D). Sequential imaging shows that an increasing number of WT nuclei (Hoechst^+^/nTom^-^) became positive for nTom during differentiation (Figure 4E-G). After 144h, when full differentiation was achieved, almost 60% of all nuclei became double positive for Hoechst and nTom (Figure 4E), a number that was similar as observed in the *in vivo* situation (single myofibers) upon exercise. This data shows that as soon as SC fuse to exisiting fibers and contribute to the myonuclear pool, during *in vitro* differentiation or *in vivo* following exercise, proteins from the newly acquired myonuclei propagate to old residing myonuclei within the same myofiber. In addition, we also observed that the original nTom^+^ myoblasts expressed higher levels of nTom when compared to the nTom levels in the differentiated cells, suggesting that nTom protein was diluted between nuclei (Figure 4F). Finally, time-lapse microscopy imaging showed that propagation occurs at very fast rates, as nTom was transported towards neighboring nTom^-^ myonuclei within 15 min of myoblast fusion (Figure 4-video supplement 1). In conclusion, it is not possible to determine the exact number of SCs that contribute to myonuclei using common lineage tracing reporter models.

**Figure 4.**
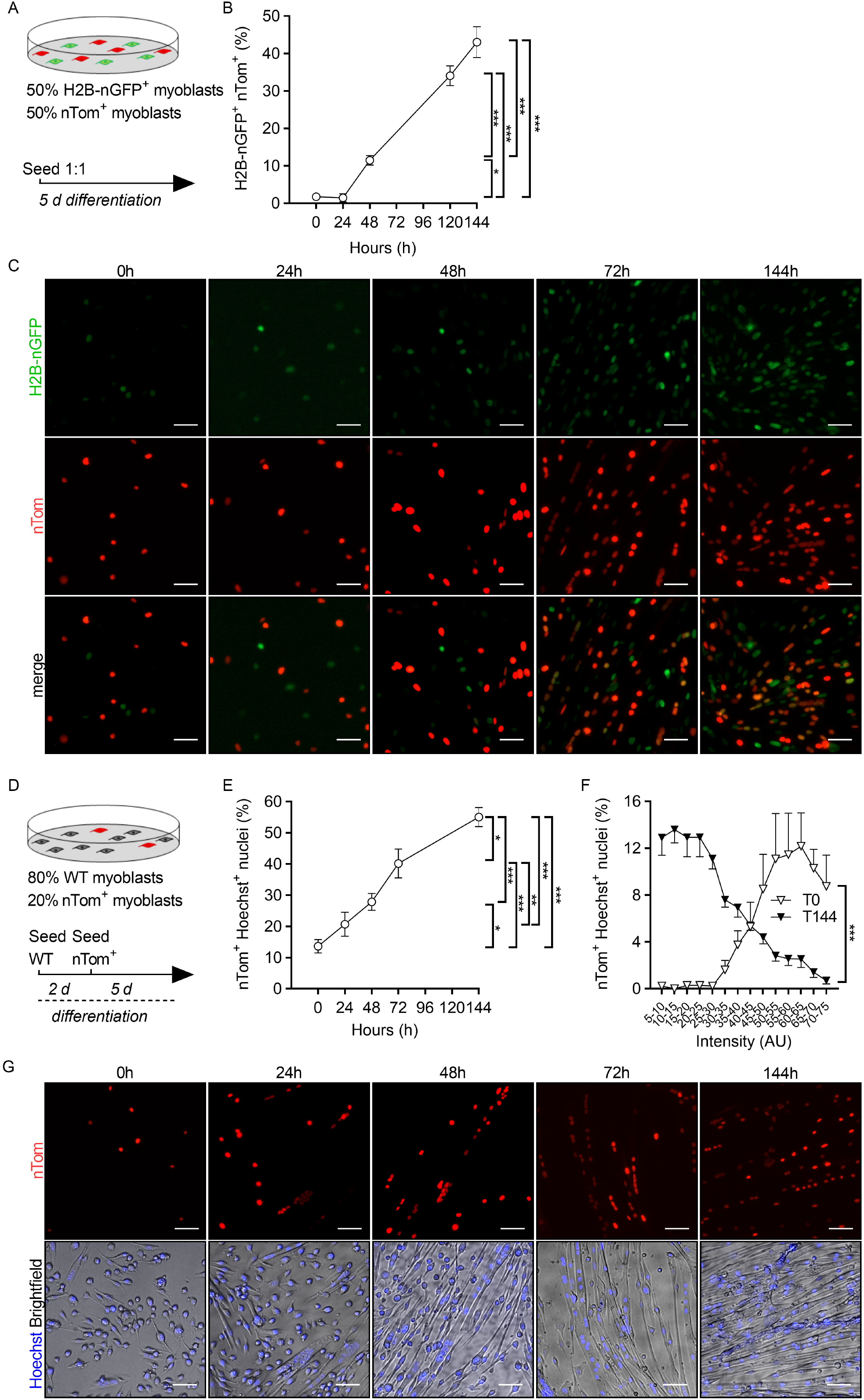
Fuorescent nuclear protein propagation renders reporter mice inadequate to assess SC fusion in myofibers. (A) Nuclear H2B-nGFP expressing and nTom^+^ myoblast were seeded at 1:1 ratio and differentiated for 144h. (B) Double positive nuclei for H2B-nGFP^+^ and nTom^+^ (marked in yellow) were assessed over time in relation to total nuclei (Hoechst^+^). (C) Representative pictures of H2B-nGFP^+^ and nTom^+^ myoblast co-cultures over time. (D) nTom^+^ and committed WT myoblast were seeded at a 1:4 ratio and differentiated for 144h. (E) Double positive nuclei for nTom^+^ and Hoechst^+^ were assessed over time in relation to total nuclei (Hoechst^+^). (F) nTom intensity was measured 1h after seeding (T0) and upon full differentiation (T144). Data is represented as percentage of nTom^+^ nuclei per intensity range in relation to the total nTom^+^ nuclei. (G) Representative pictures of WT and nTom^+^ myoblast co-cultures over time. Statistics: one-way ANOVA test with Tukey correction for multiple comparisons (*p <0.05; **p < 0.01; ***p < 0.001). Bar graphs represent mean ± SEM (error bars). Scale bar, 50 μm.

### SC fate tracing at the DNA level confirms load-dependent SC fusion during exercise

Since our data shows that counting the number of nTom^+^ nuclei in *Pax7^nTom/+^* mice does not allow to evaluate the exact contribution of SCs to myonuclei during exercise, we used a RT-PCR on genomic DNA to measure Cre-mediated recombination. This allows to trace the fate of SC at the DNA level. To this end, we sorted Hoechst^+^/PCM1^+^ myonuclei from *Pax7^nTom/+^* Control, VRun and VResRun mice (SOL muscle)(Figure 5B) and isolated DNA from these myonuclei. We designed primers specific for the recombined sequence and an internal control (housekeeping gene) (Figure 5A). We found that after 8 weeks of VResRun, almost 20% of all myonuclei were recombined, while only 12% and 4% were recombined in VRun and Control respectively (Figure 5C). To further substantiate these data, we extracted genomic DNA from bulk muscle (SOL) from Control, VRun and VResRun *Pax7^mTmG/+^* mice and and observed a similar load-dependent increase in SC contribution to nuclei upon exercise (Figure 5D,E). Collectively, these observations show that mechanical load during exercise independently promotes SC contribution to existing myofibers which manifests in myonuclear accretion.

## Discussion

Satellite cells are required for muscle regeneration and in addition contribute to muscle homeostasis under sedentary conditions (Keefe et al., 2015; Pawlikowski et al., 2015), but how they contribute to training adaptations is still poorly understood. Using two SC fate tracing mouse models, we show that exercise promotes the contribution of SCs to myofibers in a load-dependant manner. We further demonstrate that mice which ran against an external load (VResRun) showed higher SC contribution when compared to free runners (VRun) and that SC fusion occurs to a similar extent in oxidative and glycolytic muscle fibers. Furthermore, our data shows that increased SC fusion can occur independent of muscle hypertrophy, as VRun extensively increased SC fusion without stimulation of fiber hypertrophy. Thus, in this report, we provide direct evidence that exercise-mediated SC fusion occurs in a load-dependent manner.

Recent reports have indicated that there is a substantial contribution of SC to myofibers, even in uninjured muscle (Keefe et al., 2015; Pawlikowski et al., 2015). However, the potential signals promoting SC-to-myonuclear contribution remain elusive and have been hypothesized to be both local and systemic. When rodents walk or run, most of the work is performed by the plantar flexors. The dorsi flexors undergo much less mechanical load since they only keep the foot up in the swing phase (Carlson-Kuhta et al., 1998). In this respect, recent work from our group has shown that acute VResRun increases mTORC1 signaling exclusively in SOL, PLT and the oxidative part of the GAS, but not in TA (D’Hulst et al., 2019). Moreover, muscle hypertrophy is predominantly observed in plantar flexors upon resistance running, as opposed to dorsi flexors (Call et al., 2010; Konhilas et al., 2005; Legerlotz et al., 2008; White et al., 2016). Using *Pax7^CreERT2/CreERT2^; Rosa^mTmG/mTmG^* mice, we show that voluntary running strongly increases SC fusion in the plantar flexors SOL and PLT, partially in the GAS, and not in EDL or TA dorsiflexors. Furthermore, adding resistance to the wheel further increased SC fusion, indicating that the extra muscular work independently promotes SC fusion. Thus, our data shows that local rather than systemic signals regulate exercise-induced SC fusion.

Whether SCs are required for muscle hypertrophy is a heavily debated topic (Egner et al., 2017) and likely depends on age, genetic mutation, models used to induce hypertrophy and time of sampling after the applied stimulus (Cornelison, 2018; Goh et al., 2019; Goh and Millay, 2017; Mccarthy et al., 2017; Murach et al., 2018; O’Connor et al., 2007). Moreover, impairing SC contribution by the frequently used *Pax7^CreER-DTA^* model might evoke undesirable adaptations such as impairments of proprioception and reduced voluntary wheel running, which do not reflect physiological circumstances (Jackson et al., 2015). Although we could not causally link SC fusion with hypertrophy in this study since we used reporter models to trace SC contributions under physiological circumstances, our tracing approach allows us to evaluate whether SC fusion occurs upon exercise and whether concomitant myonuclear accretion occurs in the absence or presence of hypertrophy. VResRun induced a 23% increase in SOL wet weight, which was exclusively caused by the increase in CSA of MyHCIIa fibers, while no hypertrophy was observed in VRun in any fiber type. This is remarkable given that even though SC fusion was higher in VResRun, we observed similar numbers of GFP positive MyHCI versus MyHCIIa fibers in both VRun and VResrun. Thus, the lack of hypertrophy in the heavily SC fused MyHCI fibers promotes the hypothesis that myonuclear accretion might also play a role in endurance adaptations to interval training (Abreu et al., 2017; Hawley et al., 2014) and that myonuclear accretion is not merely a way to withstand to expansion of the myonuclear domain (Conceiçao et al., 2018).

The *Pax7^mTmG/+^* mice only provide a rough estimation of SC contribution to myonuclei under exercise conditions since the mGFP diffuses throughout the sarcolemma. Therefore, to get a quantitative insight into the extent of SC contribution to myonuclei, we used a nuclear reporter mouse in which the fluorescent reporter (nTom) is retained exclusively within the Pax7^+^ cells and all SC-derived myonuclei. Strikingly, we found that nearly all myonuclei became nTom^+^ after 8 weeks of VRun and VResRun, despite only a modest 14% and 16% increase in myonuclear number by VRun and VResRun respectively. During the preparation of this manuscript, an extensive report provided direct evidence of fluorescent protein traveling between nuclei of primary differentiated myotubes, a phenomenon termed nuclear propagation (Taylor-weiner et al., 2020). In concordance, spatial distribution of specific transcription factors, which may exit the myonucleus and activate receptors in other regions of the myofiber has been reported (Fontaine et al., 1988), suggesting nuclear transport of various proteins between nuclei in myofibers. We confirmed propagation in our model by a series of co-culturing experiments using H2B-nGFP and nTom expressing myoblasts. After 6 days of differentiation, when myotubes reached full differentiation, nearly 60% of all nuclei became nTom^+^, while WT and nTom were initially seeded at a 1 to 4 ratio. Similar results were obtained when nTom^+^ and H2B-nGFP^+^ myoblasts were differentiated in the same culture. Thus, the high number of nTom^+^ myonuclei, which we observed after voluntary running is a vast overestimation of SC fusion. Importantly, these data warrant against the use of nuclear reporter mice in multinucleated cells such as myofibers and suggest a careful re-interpretation of previous data published with *Pax7* nuclear reporter mice (Bachman et al., 2018; Liu et al., 2017; Tierney et al., 2017).

To circumvent nuclear propagation artefacts in our *Pax7^nTom/+^* mice, we assessed recombination at the DNA level from both sorted myonuclei as well as bulk muscle derived from two *Pax7* reporter models. Intriguingly, the increase in the fraction of recombined myonuclei from sedentary control to VRun (+8%) and (VResRun) (+12%) was almost identical to the increases in myonuclear accretion measured on single fiber (Hoechst^+^) and on cryosections (PCM1^+^). Myonuclei have a half-life of at least 15-years (Spalding et al., 2005) and are remarkably resistant to apoptosis, even when the myofiber is challenged by unloading or atrophy (Bruusgaard and Gundersen, 2008). This suggests that myonuclear turnover during exercise training is low, and that the ~16% increase in myonuclei upon exercise as observed in this study is the direct result of SC fusion to myofibers, without active nuclear apoptosis. Thus, based on our data, a major question that arises is whether these new myonuclei are different in terms of transcriptional or epigenetic signature from the old residing ones. In addition, whether they contribute to training adaptations and whether there is a difference between endurance and resistance-based training is yet to be explored. Future research using single nuclei transcriptomics may elucidate potential reprogramming of newly derived myonuclei during exercise.

In summary, this study is the first to assess SC fusion and concomitant myonuclear accretion in a physiological setting of increased mechanical load. Using a *Pax7* membrane-marker reporter, we show increased load-dependent SC fusion in the plantar flexors of the mouse hindlimb after 8 weeks of voluntary running. Predominantly the glycolytic type MyHCIIa fibers in SOL underwent hypertrophy, but SC fusion occurred to the same extent in both in MyHCI and MyHCIIa fibers. Thus, the lack of hypertrophy in the heavily SC fused MyHCI fibers strongly suggests that myonuclear accretion might also play a role in endurance adaptations to interval training. Furthermore, in an attempt to objectively quantify SC-derived myonuclear accretion after voluntary running, we used a *Pax7* nuclear reporter mouse and unexpectedly observed massive increase in nTom positive nuclei. Further *in vitro* coculture experiments showed that this was due to propagation of nTom between nuclei sharing the same cytoplasm. Hence, future investigations should warrant caution in using nuclear reporter constructs in multinucleated cells. Finally, assessment of Cre-mediated recombination in *Pax7^CreERT2^* reporter models indicated that SC contribution to myofibers mirrors myonuclear accretion during exercise.

## Acknowledgements

We gratefully acknowledge Nicola Bundschuh and Paola Gilardoni for their skilled technical assistance and Andrew Palmer for his thoughtful discussions about the experiments.

## Author contributions

Evi Masschelein; conceptualization, data curation, formal analysis, investigation, methodology, writing original draft, review and editing, Gommaar D’Hulst; conceptualization, data curation, formal analysis, investigation, methodology, writing original draft, review and editing, Laura Hinte; formal analysis, methodology, manuscript editing, Joel Zvick; formal analysis, methodology, manuscript editing, Inés Soro-Arnaiz; formal analysis, methodology, manuscript editing, Tatiane Gorski; formal analysis, methodology, manuscript editing, Ferdinand von Meyenn: conceptualization, investigation, manuscript editing Ori Bar-Nur; conceptualization, investigation, manuscript editing and Katrien De Bock; conceptualization, formal analysis, funding acquisition, investigation, methodology, writing original draft, project administration and editing.

---

**Figure 1-figure supplement 1. Running pattern**. (A) Average running speed, (B) running time and (C) number of bouts per night throughout the exercise protocol (n = 16 mice per group). Statistics: two-way ANOVA test with a Bonferroni post hoc test. Group x time interaction is indicated in the figure. (*p <0.05; **p < 0.01; ***p < 0.001). Line graphs represent mean ± SEM (error bars).

**Figure 1-figure supplement 2. Muscle hypertrophy is muscle dependent, fiber type dependent and load-dependent**. (A) Representative cross-sections for m. soleus (SOL) stained for MyHCI (green), MyHCIIa (blue) and Laminin (red). (B,C) Quantification of fiber cross-sectional area distribution and (D,E) average fiber crosssectional area. (F) Representative cross-sections for plantaris (PLT) stained for MyHCIIa (blue), MyHCIIb (yellow) and laminin. (G,H) quantification of fiber crosssectional area distribution and (I,J) average fiber cross-sectional area. (K) Fiber cross-sectional area distribution for gastrocnemius (GAS), (L) tibialis anterior (TA) and (M) extensor digitorum longus (EDL). Statistics: one-way ANOVA test with Tukey correction for multiple comparisons (*p < 0.05; **p < 0.01; ***p < 0.001) (a, p < 0.05 VResRun compared to Control; b, p < 0.05 VResRun compared to VRun; c, p < 0.05 VRun compared to Control). Bar graphs and line graphs represent mean ± SEM (error bars). Scale bar, 100 μm.

**Figure 2-figure supplement 3. SC fusion is not fiber type dependent.** (A) Representative sequential cross-sections for soleus for mGFP and mTOMATO (left panel) and stained for MyHCI (yellow) and MyHCIIa (blue) (right panel). (B) Quantification of GFP^+^ fibers for MyHCI and (C) MyHCIIa. (D) Fiber type distribution and (E) fiber type distribution of GFP^+^ fibers. Statistics: one-way ANOVA test with Tukey correction for multiple comparisons (*p < 0.05; **p < 0.01; ***p < 0.001). Each dot represents a single mouse. Bar graphs and line graphs represent mean ± SEM (error bars). Scale bar, 100 μm.

**Figure 3-figure supplement 4. Myonuclear accretion is load-dependent.** (A) Representative cross-sections of soleus stained for PCM1 (yellow), Hoechst (blue) and wheat germ agglutinin (WGA) (red). (B) Quantification of PCM1^+^ nuclei per fiber. Statistics: one-way ANOVA test with Tukey correction for multiple comparisons (*p < 0.05; **p < 0.01; ***p < 0.001). Bar graphs and line graphs represent mean ± SEM (error bars). Scale bar, 100 μm.

**Figure 4-video supplement 1. In vitro nuclear propagation.** Time-lapse epifluorescence microscopy of a fusing nTom^+^ myoblast with WT myoblasts (nTom^-^). White arrow shows propagation of nTom. Scale bar, 50 μm.

**Figure 1-table supplement 1.** Phenotypic characterization and muscle weights normalized to tibia length. Statistics: one-way ANOVA test with Tukey correction for multiple comparisons (*p < 0.05 VRun/VResRun vs. Control.). Values represent mean ± SEM. n = 9-12 mice per group.

|  | Control | VRun | VResRun |
| --- | --- | --- | --- |
| **Whole muscle weights** |  |  |  |
| SOL (mg)/tibia length (cm) | 5.15. ± 0.28 | 5.38 ± 0.21* | 6.39 ± 0.36* |
| PLT (mg)/tibia length (cm) | 7.10 ± 0.39 | 7.39 ± 0.21 | 7.13 ± 0.44 |
| GAS (mg)/tibia length (cm) | 58.81 ± 3.50 | 58.74 ± 1.09 | 56.70 ± 1.65 |
| TA (mg)/tibia length (cm) | 24.04 ± 0.95 | 23.79 ± 0.72 | 21.39 ± 0.91 |
| EDL (mg)/tibia length (cm) | 5.53 ± 0.29 | 5.49 ± 0.15 | 5.24 ± 0.20 |
| **Phenotype** |  |  |  |
| Body mass (g) | 24.02 ± 0.51 | 24.23 ± 0.49 | 24.26 ± 0.65 |
| Lean mass (g) | 20.76 ± 0.53 | 20.67 ± 0.43 | 21.12 ± 0.57 |
| Fat mass (g) | 2.35 ± 0.12 | 2.32 ± 0.15 | 2.56 ± 0.21 |

