## Supplementary figures and images for "Exercise promotes satellite cell contribution to myofibers in a load-dependent manner"

### Supplemental Fig 1

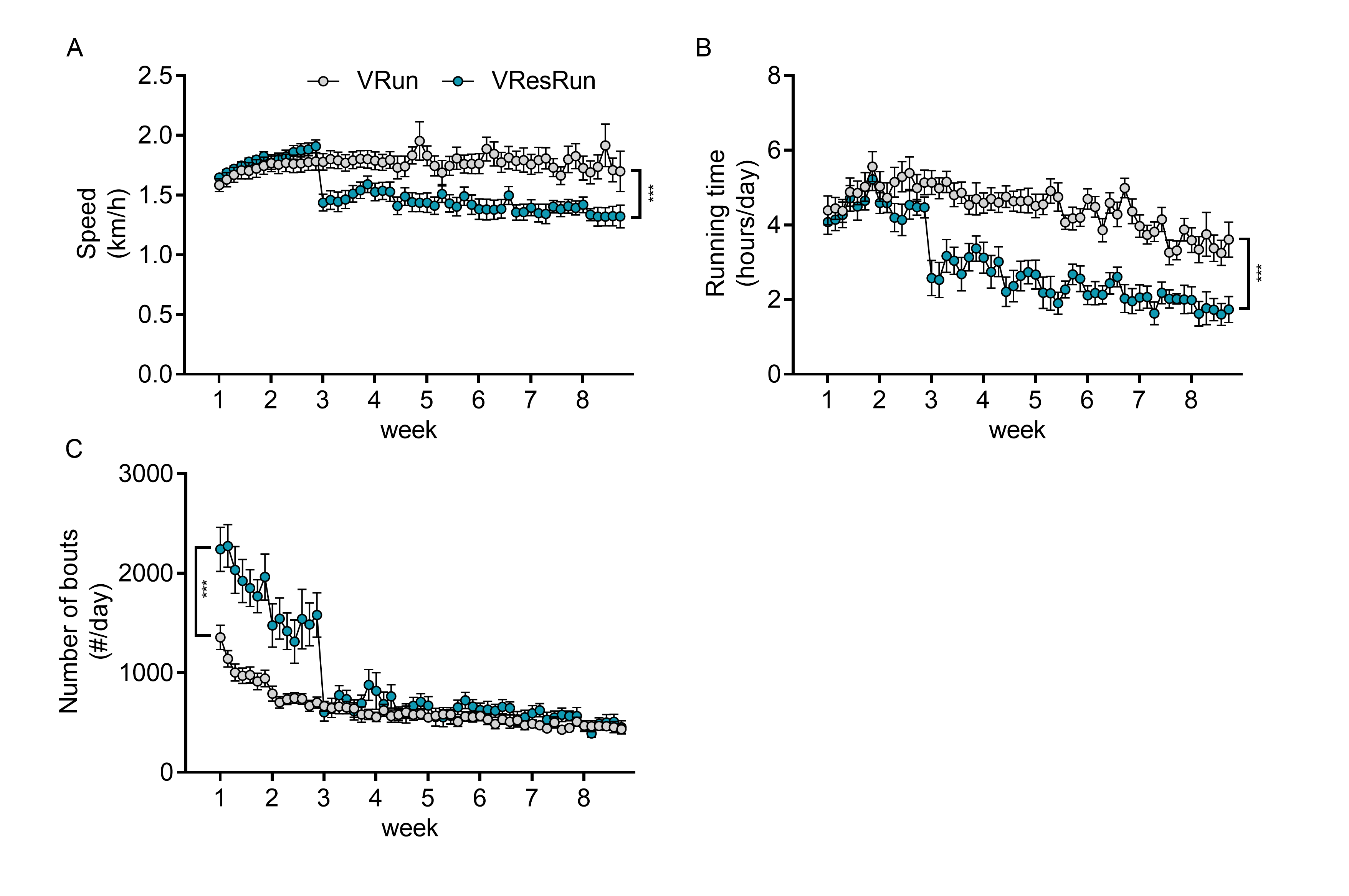

### Supplemental Fig 2

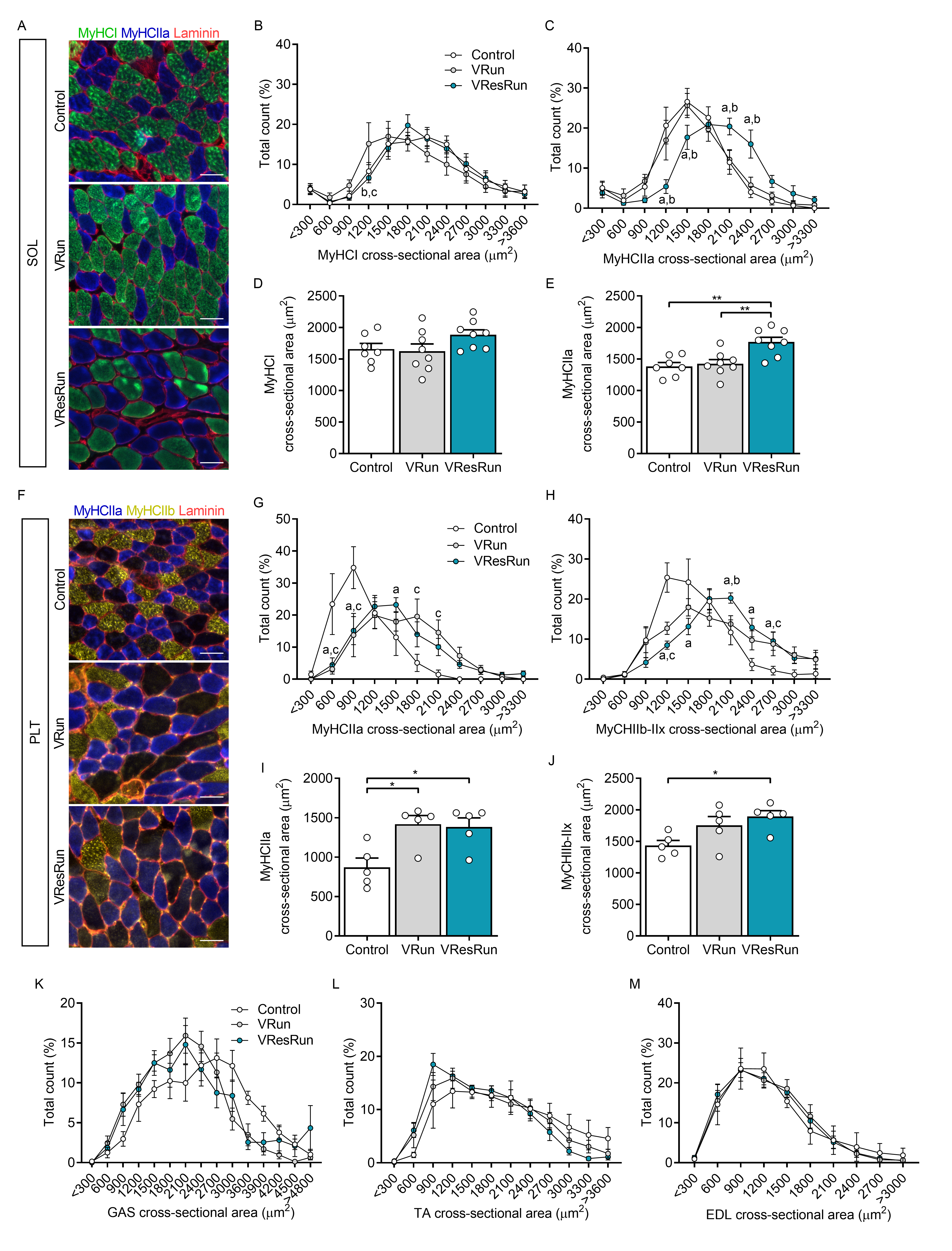

### Supplemental Fig 3

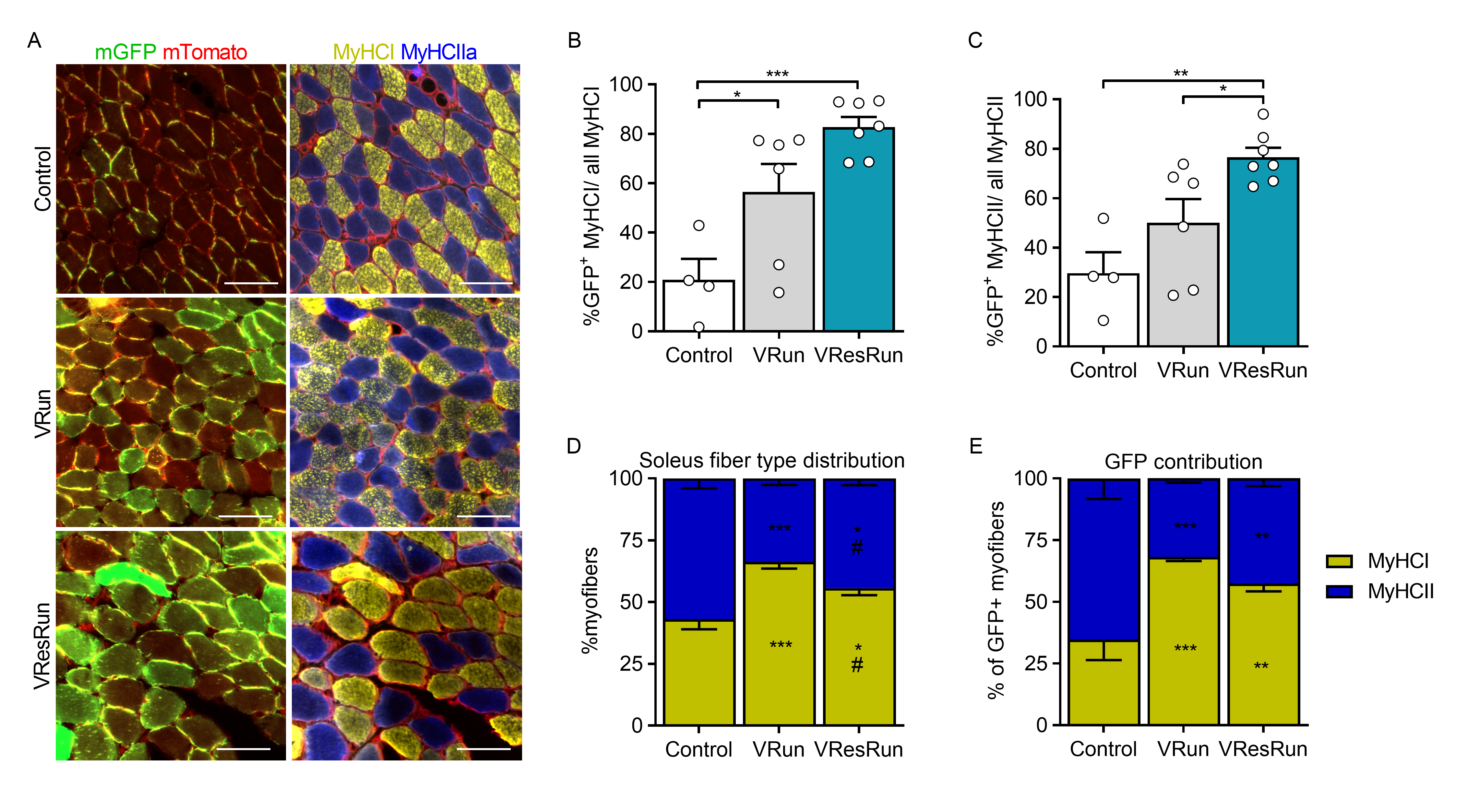

### Supplemental Fig 4

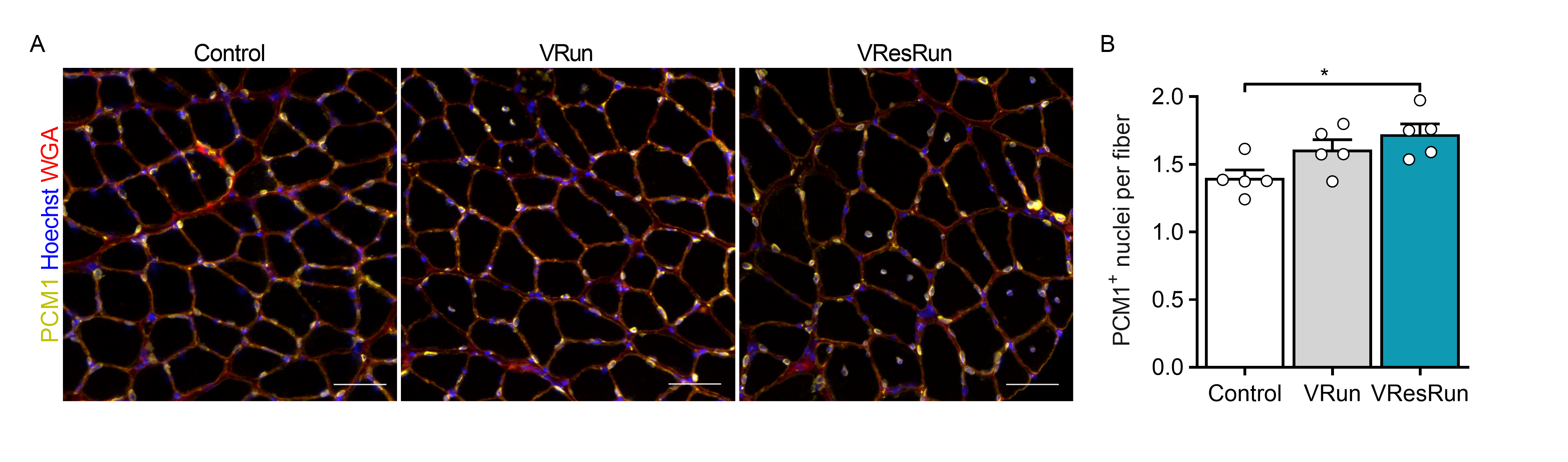
